# RNA-GPS Predicts SARS-CoV-2 RNA Localization to Host Mitochondria and Nucleolus

**DOI:** 10.1101/2020.04.28.065201

**Authors:** Kevin Wu, James Zou, Howard Y. Chang

## Abstract

The SARS-CoV-2 coronavirus is driving a global pandemic, but its biological mechanisms are less well understood. SARS-CoV-2 is an RNA virus whose multiple genomic and sub-genomic RNA (sgRNA) transcripts hijack the host cell’s machinery, located across distinct cytotopic locations. Subcellular localization of its viral RNA could play important roles in viral replication and host antiviral immune response. Here we perform computational modeling of SARS-CoV-2 viral RNA localization across eight subcellular neighborhoods. We compare hundreds of SARS-CoV-2 genomes to the human transcriptome and other coronaviruses and perform systematic sub-sequence analyses to identify the responsible signals. Using state-of-the-art machine learning models, we predict that the SARS-CoV-2 RNA genome and all sgRNAs are enriched in the host mitochondrial matrix and nucleolus. The 5’ and 3’ viral untranslated regions possess the strongest and most distinct localization signals. We discuss the mitochondrial localization signal in relation to the formation of double-membrane vesicles, a critical stage in the coronavirus life cycle. Our computational analysis serves as a hypothesis generation tool to suggest models for SARS-CoV-2 biology and inform experimental efforts to combat the virus.

## Introduction

COVID-19 (coronavirus disease 2019) has become a global pandemic, fueled by the rapid spread of the coronavirus SARS-CoV-2 (severe acute respiratory syndrome coronavirus 2), a positive strand RNA virus (Sanche et al., 2020; Wu et al., 2020a). The scientific community is actively trying to understand SARS-CoV-2’s biological mechanisms and effects. Here, we computationally analyze the subcellular localization patterns of SARS-CoV-2 RNA transcripts. Our results suggest potential avenues for experimental validation and follow-up, while providing a template for *in silico* analyses of viral biology.

RNA subcellular localization is critical to a myriad of cellular processes (Buxbaum et al., 2015; Chin and Lécuyer, 2017; Ryder and Lerit, 2018). Researchers have also discovered that RNA localization plays a significant role in RNA viruses (Chou et al., 2013), with functions ranging from regulating sites of virion assembly (Becker and Sherer, 2017) to disrupting host mitochondrial function (Somasundaran et al., 1994). Thus, understanding the behavior and localization of SARS-CoV-2 RNA transcripts can lead to a better understanding of its function and pathogenicity, potentially revealing targetable mechanisms.

We perform computational modelling for SARS-CoV-2 subcellular RNA localization. In particular, we build upon our recent work developing RNA-GPS, a state-of-the-art computational model predicting high-resolution RNA localization in human cells (Wu et al., 2020b), trained on transcriptome-wide localization patterns of human RNAs across eight subcellular landmarks (Fazal et al., 2019). RNA-GPS’s strong performance, coupled with viruses’ dependence on hijacking existing cell machinery for reproduction, suggests that RNA-GPS could provide insights into SARS-CoV-2’s localization behavior and can focus future experimental efforts.

We use RNA-GPS to interrogate the localization patterns of SARS-CoV-2’s genome, which spans approximately 30 kilobases of single-stranded positive-sense RNA (Kim et al., 2020) (Figure 1a). We predict that SARS-CoV-2 is enriched for localization in the nucleolus and the mitochondria. Comparison of SARS-CoV-2’s predicted localization against that of other human coronaviruses, including strains causing the common cold, Middle East respiratory syndrome (MERS), and the SARS outbreak of 2003, shows that SARS-CoV-2 exhibits a stronger mitochondrial and nuclear localization signal than a large majority of its coronavirus relatives. We additionally find that this localization signal appears to be driven by many redundant motifs spread across the genome, suggesting its importance. We conclude by connecting our observations to known RNA and viral biology, proposing possible explanatory mechanisms for previously observed phenomena. Our analysis suggests experimental validation of our predictions and serves as a framework for applying data science for principled hypothesis generation, enabling targeted response to this and future outbreaks.

**Figure 1:**
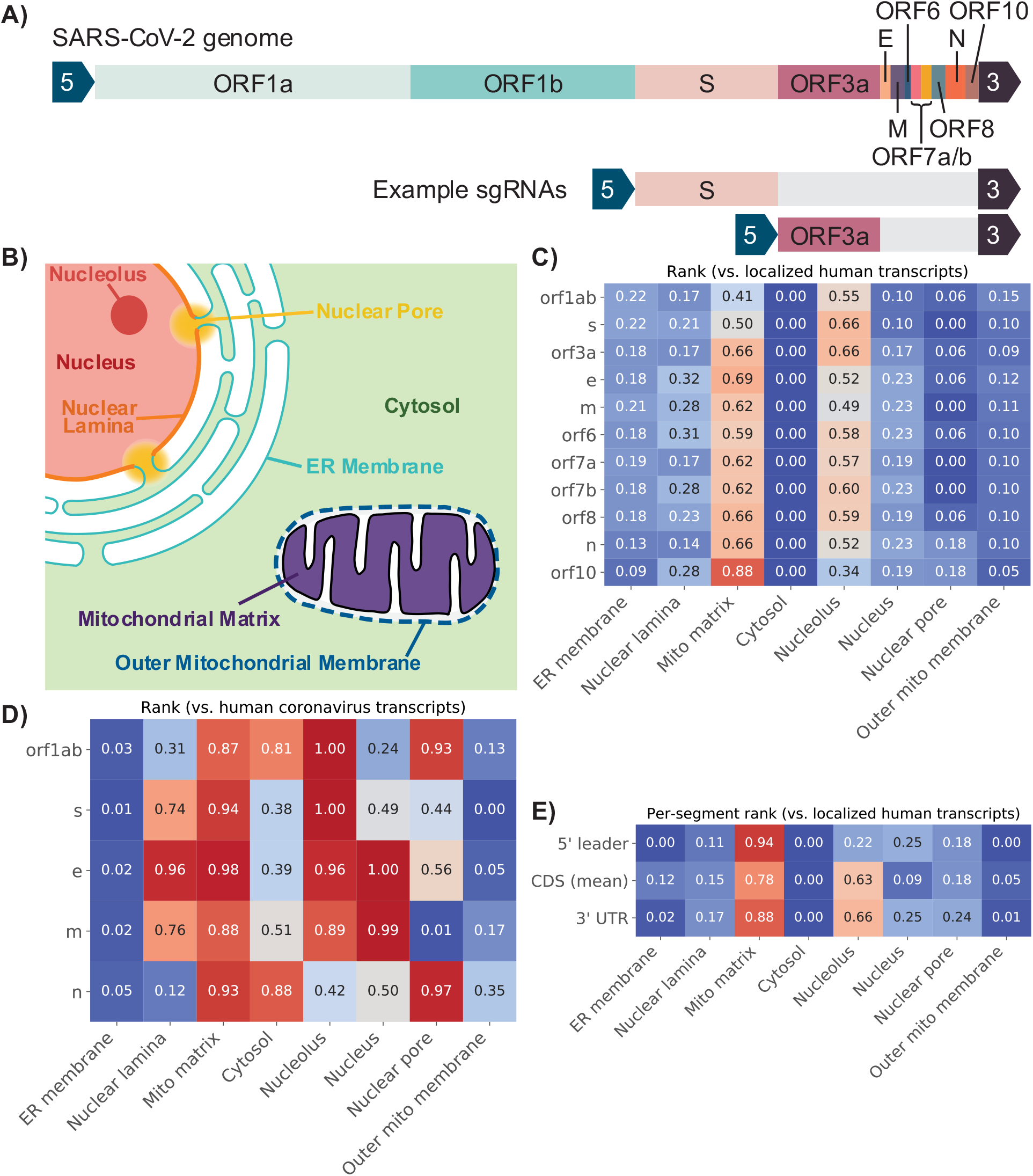
Depictions of the SARS-CoV-2 genome (a), the eight compartments that RNA-GPS predicts transcript localization to (b), and the predicted localizations for SARS-CoV-2 sgRNAs (c, d) and its individual 5’/CDS/3’ sequence segments (e). The SARS-CoV-2 genome produces a series of sub-genomic RNAs (sgRNAs), each encoding one or more genes/proteins (a). These sgRNAs share a common leader 5’ sequence and a common trailing 3’ UTR sequence (arrow blocks). For each sgRNA, RNA-GPS predicts localization to each compartment shown in (b) (figure reproduced from (Wu et al., 2020b)), the results of which are shown in (c). This heatmap shows rank scores, indicating how strongly each sgRNA (rows) localizes to each compartment (columns), compared to a typical endogenous human transcript localizing to that compartment. Colors directly correlate with indicated rank scores. Most sgRNAs exhibit similar localization patterns, with a general enrichment towards the mitochondrial matrix and nucleolus. We also computed these rank scores against a baseline of other coronavirus localization signals in (d). SARS-CoV-2 exhibits a stronger mitochondrial matrix localization signal than most other coronaviruses, along with greater overall nuclear localization, particularly at the nucleolus. Interestingly, localization to the ER membrane is not present as a particularly strong signal in either context. (e) Shows the predicted localization rank scores for shared 5’ and 3’ segments, and an averaged localization rank score for CDS segments. Even on their own, the short ~90-250 base pair 5’ and 3’ segments carry the mitochondrial and nucleolar localization signals.

## Results

We leverage our recent work developing RNA-GPS, a computational model predicting high-resolution RNA localization in human cells trained with HEK293T APEX-seq data (Wu et al., 2020b). Briefly, RNA-GPS predicts localization of RNA transcripts to eight different subcellular locations: the cytosol, endoplasmic reticulum, mitochondrial matrix, outer mitochondrial membrane, nucleus, nucleolus, nuclear lamina, and nuclear pore (Figure 1b), and has been shown to generalize well to additional cell lines including HeLa-S3 and K562. Although RNA-GPS is trained on human, not viral, RNA transcripts, its strong test performance combined with the fact that viruses commandeer human cellular machinery suggest that it offers a reasonable hypothesis of viral transcript localization behavior given currently available data.

We consider average localization predictions to each compartment across all released and annotated SARS-CoV-2 genomes available as of April 6, 2020 (n = 213) on GenBank (Coordinators, 2018). SARS-CoV-2 is believed to enter the cell as a positive strand genomic RNA, subsequently forming 11 positive strand sub-genomic RNA (sgRNA) transcripts encoding different open reading frames and sharing the same 5’ leader sequence and 3’ untranslated region (UTR) (Figure 1a). Within each viral genome, we predict the localization of each sgRNA produced from the primary SARS-CoV-2 genome. To provide more context, we frame these predicted localization probabilities relative to the predictions of other relevant “baseline” transcript sequences. We consider two such baselines: the distribution of model predictions on transcripts exhibiting significant enrichment within the human HEK293T cell line (n = 366 transcripts) (Fazal et al., 2019), and the distribution of model predictions on transcripts derived from human coronaviruses, excluding SARS-CoV-2 (n = 191 genomes, spanning diseases from the common cold to MERS, Supplementary Figure 1a). The human baseline gives us an idea of how strong localization signals within SARS-CoV-2 are, relative to naturally occurring human transcripts with well-characterized localization behaviors. The coronavirus baseline focuses on differences in the localization behavior of SARS-CoV-2 relative to similar viral specimens – differences that may help researchers focus on the particularities of this virus. For both baselines, we calculate the proportion of the baseline distribution that the SARS-CoV-2 localization prediction exceeds, which we refer to as a rank score. For example, a localization score of 0.6 for the nucleolus relative to human transcripts suggests that the particular viral RNA is more likely to localize to the nucleolus compared to 60% of the human RNAs that shows some localization to the nucleolus.

### SARS-CoV-2 RNA subcellular localization patterns

We find that compared to transcripts with known localizations in human cells, SARS-CoV-2 has a notable localization signal towards the mitochondrial matrix, as well as the nucleolus (Figure 1c). We observe consistent localization predictions of different sgRNAs encoded by the virus (shown in each row, Figure 1c). Interestingly, we do not see particularly strong transcript localization towards the ER membrane or the cytosol, compartments canonically known to harbor viral transcripts and proteins for replication and assembly. However, prior works have shown that some RNA viruses exhibit transcript localization to mitochondria (Somasundaran et al., 1994), and that the nucleolus plays a prominent role in the viral life cycle, even for viruses that primarily replicate in the cytoplasm as SARS-CoV-2 does (Salvetti and Greco, 2014).

In addition to framing our results with respect to endogenous human transcripts, we also compare predicted localization signals of SARS-CoV-2 sgRNAs to that of other human coronaviruses (Figure 1d). Here, we observe similar overall trends in our localization predictions. We see that the mitochondrial matrix localization signal previously described is recapitulated here, suggesting that not only does SARS-CoV-2 have a mitochondrial matrix localization signature comparable to that of human transcripts also localizing to the mitochondrial matrix, but this signal is stronger than that of other coronaviruses. Additionally, we see an overall pattern suggesting that SARS-CoV-2 may have a greater affinity for nuclear localizations (nuclear pore, nucleus, nucleolus, and nuclear lamina) compared to other coronaviruses. Similar to our prior results, we also see that the ER membrane localization signals are weaker for SARS-CoV-2, even compared to other human coronaviruses.

We also evaluated how the viruses comprising the coronavirus baseline themselves localize, compared to human transcripts. We found that the most prominent localization signals for general human coronaviruses pointed towards the nucleolus, mitochondrial matrix, and ER membrane (Supplementary Figure 1b). Overall, our computational analysis suggests that SARS-CoV-2’s sgRNA transcript localization towards the mitochondrial matrix and nucleolus may be amplifications of localization behaviors that were already present in coronaviruses.

To validate the robustness of these localization trends, we also trained a different predictive algorithm (a recurrent neural network, see Methods section for additional details) on the APEX-seq data and performed a similar set of experiments, comparing SARS-CoV-2 localization predictions to human and coronavirus baselines. This alternative model also predicts strong mitochondrial matrix and nucleolus localization for SARS-CoV-2 (Supplementary Figure 2). Since this algorithm uses a very different modeling strategy as RNA-GPS and converges to similar findings, this suggests that the mitochondrial matrix and nucleolus signals are not artifacts of a particular model and increases our confidence in the findings.

### SARS-CoV-2 negative strand RNA also localizes to mitochondria and nucleolus

During its replication life cycle, SARS-CoV-2 copies its positive strand RNA to create a negative strand RNA that serves as the template for viral “transcription” and production of sgRNAs (Wu and Brian, 2010). We applied RNA-GPS to the negative strand RNA and discovered that the negative strand RNA also exhibits localization signal to the mitochondrial matrix and nucleolus, albeit weaker than the positive strand sequence (Supplementary Figure 3). This result suggests that the sequence features driving these localization patterns are independently present in both positive and negative strand RNAs and can potentially drive viral double-stranded RNA accumulation at specific subcellular locales.

### SARS-CoV-2 5’ and 3’ UTRs contain strong localization signals

In addition to predicting localization, our computational model can also help understand which regions of the transcript may be more responsible for driving these observed localizations. We specifically investigated the potential contribution of the three main regions of the SARS-CoV-2 genome: the shared 5’ leader sequence, the shared 3’ UTR, and the variable “coding” sequence in the middle. We predicted the localization for each of these regions by itself (Figure 1e). The 5’ leader sequence shows the strongest localization signal to the mitochondrial matrix, and no signal for the nucleolus. In contrast, the 3’ UTR has the strongest localization for the nucleolus and also has strong signal for the mitochondrial matrix. The coding sequence (CDS) also shows specific signals for these two compartments. Because the 5’ and 3’ sequences are shared by the different SARS-CoV-2 sgRNAs, this is likely a strong factor behind the consistent localization patterns we find across the different sgRNAs. We performed further computational ablation studies of RNA binding protein (RBP) motifs in SARS-CoV-2 and found that the known motifs do not appear to be the primary drivers of the mitochondrial matrix and nucleolus localization patterns. Specifically, computational deletions of all instances of each RBP motif in SARS-CoV-2 sequence, repeated across all enriched RBPs, did not significantly alter the RNA-GPS predictions. This result suggests that the SARS-CoV-2 localization signal is highly redundant within the viral genome.

## Discussion

In this work, we apply computational models of human RNA transcript localization to better understand the subcellular localization of the SARS-CoV-2 genome and its constituent sgRNAs. This approach builds upon the idea that viruses must use existing human cell machinery to reproduce, and consequently that sequence-based localization signals are likely shared between human and coronavirus transcripts. The strengths of this approach include (1) the potential to understand viral RNA localization without the risk of live viral cultures; (2) the ability to examine hundreds of viral isolates and related coronaviruses and thousands of RBP motif ablations; (3) the ability to examine viral genes, UTRs, and negative strands individually, which may otherwise require the ability to precisely synchronize and arrest the viral life cycle. We find that SARS-CoV-2 appears to harbor strong localization signals towards the mitochondrial matrix and nuclear compartments, comparable to human RNA and more so than other coronaviruses. This intriguing hypothesis suggests future experimental exploration and validation.

These results might appear surprising, as one might expect localization signals to enrich towards regions like the endoplasmic reticulum, where viral translation, viral assembly, and disruption of normal cell activity are commonly known to occur (Fung and Liu, 2014; Minakshi et al., 2009; Nal et al., 2005). However, coronaviruses are known to produce complex double-membrane vesicle (DMV) structures during viral replication, which may serve functions like concealing the virus from cellular defenses (Hagemeijer et al., 2012; Knoops et al., 2008). While these DMVs are generally believed to be formed via viruses manipulating the ER membrane (Blanchard and Roingeard, 2015), the mechanism for importing and packaging proteins and RNA into these miniature organelles is not as clearly understood. One possible mechanism for importing viral RNA involves the virus exploiting endogenous RNA localization mechanisms that the cell already possesses for existing double-membrane organelles: namely, the mitochondria. Indeed, introducing just 2 amino acid point mutations in the murine coronavirus can cause both a significant drop in the number of DMV structures observed, as well as a sharp increase in viral protein localization at the mitochondria (Clementz et al., 2008). This suggests a high degree of resemblance between the DMV and mitochondrial localization mechanisms – leading to the hypothesis that our mitochondrial matrix localization predictions are capturing this similarity between the DMV and mitochondria. Furthermore, DMVs have been shown to contain double-stranded RNA (Hagemeijer et al., 2012), and our strand-agnostic localization predictions are concordant with this evidence. Under this model, SARS-CoV-2’s strong mitochondrial localization signal relative to other coronaviruses may even contribute to its similarly high infectivity by increasing its efficacy in generating and importing materials into these DMV structures.

Another possible interpretation of these localization results is that previously studied viral protein localizations are actually driven by transcript-level localizations, a mechanism that is highly prevalent for human proteins (Blower, 2013). Protein-protein interaction studies performed on SARS-CoV-2 have found that its NSP5 (within ORF1a), NSP13 (within ORF1b), ORF6, and ORF10 proteins interact with host proteins that predominantly localize to nuclear compartments (Gordon et al., 2020). The same study found viral interactions with Tomm70, a mitochondrial import receptor that plays a critical role in modulating interferon response – a critical anti-viral cellular defense pathway (Liu et al., 2010). In both cases, localized transcripts could be driving protein localization, enabling more focused protein-protein interactions.

Additional protein localization patterns appear within SARS-CoV-2’s more thoroughly-studied relative, the SARS-CoV coronavirus, responsible for the 2003 SARS outbreak (Ksiazek et al., 2003). The nucleocapsid (N) protein has been shown to dynamically localize to the nucleolus (Cawood et al., 2007). The transcribed protein corresponding to ORF3 of SARS-CoV localizes to both the mitochondria (Yuan et al., 2006) and the nucleolus, causing cell cycle arrest and apoptosis (Yuan et al., 2005). Some even suggest a possible mechanism where translocation of this protein between the nucleus and mitochondria influences the cell’s interferon response (Freundt et al., 2009). It is plausible that this protein localization behavior is conserved across both SARS-CoV and its contemporary SARS-CoV-2, especially given their relatively high sequence homology (Lu et al., 2020), and is likewise driven by underlying transcript localization. Indeed, these patterns can also be found much more broadly across viral species. For coronaviruses in general, N protein localization to the host nucleolus appears to be a fairly conserved functional attribute and may play a role in disrupting cell division and inhibition of cytokinesis (Rawlinson and Moseley, 2015; Wurm et al., 2001). Within the influenza A virus, nonstructural protein 1 localizes to the mitochondria (Tsai et al., 2017). For many viruses that primarily replicate within cytoplasmic regions, multiple studies have found viral proteins that nonetheless localize to the nucleus to aid replication and disrupt host cell functionality (Hiscox, 2003; Weidman et al., 2003).

One limitation of our work lies within its attempt to generalize models trained on human RNA transcript localization data, to transcripts derived from a different species. A model learned on HEK293T cells also may not be an appropriate model for cell types that are infected by SARS-CoV-2. Although the sharing of biological machinery between human cells and SARS-CoV-2 coupled with RNA-GPS’s strong performance on held-out test datasets leads us to believe that this approach is promising, viral infection also substantially remodels the cell’s internal machinery, and the expression of viral RNA binding proteins (not accounted for in our model) can both introduce errors into our predictions. Thus, localization experiments are necessary to validate our computational analyses. Cross-referencing our results against existing literature is somewhat limited, as most studies have focused on the localization of viral proteins rather than viral transcripts. Furthermore, RNA localization is undoubtedly one of many pieces of complex, interconnected mechanisms that this coronavirus adopts, and our hypotheses presented here do not preclude (many) additional critical biological phenomena.

In summary, we build upon recent computational models of RNA subcellular localization to study, *in silico,* the localization properties of SARS-CoV-2. Our results suggest that nuclear-mitochondrial transcript localization patterns may be an important, unique characteristic of SARS-CoV-2 that warrants additional study. We connect these observations to known viral biology regarding DMV structures in viral replication, as well as known protein localization patterns. In doing so, we propose potential cellular mechanisms that underpin viral biology - mechanisms that warrant experiments validating their accuracy, and perhaps even their potential as therapeutic targets. More broadly, we hope that our study helps define a framework for applying machine learning models to enable focused hypothesis generation, inspiring similar studies that leverage data science to rapidly respond to emerging epidemiological challenges.

## Code and data availability

All data used to train RNA-GPS and auxiliary models is available through the Gene Expression Omnibus under accession GSE116008. All SARS-CoV-2 genomes analyzed are publicly available through the National Center for Biotechnology Information GenBank database. Code to query for these sequences and perform downstream analysis is available at https://github.com/wukevin/rnagps, specifically under the “covid19” directory.

## Acknowledgements

We thank the members of the Chang and Zou laboratories for helpful discussions. Kevin R. Parker and Furqan M. Fazal provided particularly helpful feedback regarding biological interpretation of localization results. H.Y.C. is supported by RM1-HG007735 and R01-HG004361. H.Y.C. is an Investigator of the Howard Hughes Medical Institute. J.Z. is supported by NSF CCF 1763191, NIH R21 MD012867-01, NIH P30AG059307, and grants from the Silicon Valley Foundation and the Chan-Zuckerberg Initiative.

## Author Contributions

H.C. and J.Z. conceived the idea for this project and supervised its execution. K.W. gathered, preprocessed, and analyzed data for this project with input from all authors. K.W. wrote the manuscript with significant input from all authors.

## Methods

### Obtaining viral genomes

SARS-CoV-2 viral genomes were programmatically queried from the GenBank online database using the BioPython library’s Entrez module (Cock et al., 2009). Returned results were then filtered to retain only assemblies that included annotated, named sgRNA “genes.” In cases where the shared 5’ leader sequence or the 3’ tail were not explicitly annotated, their regions were inferred to be the 5’ and 3’ trailing bases outside of any coding regions, respectively. As there are many SARS-CoV-2 genome assemblies that fit these criteria, localization predictions are averaged across all genomes.

Viral genomes constituting the coronavirus baseline follow an identical process, save for using a different query sequence with GenBank that specifically fetches matches to the six known coronaviruses known to infect humans (excluding SARS-CoV-2): 229E, NL63, OC43, HKU1, MERS-CoV (beta coronavirus that causes Middle East Respiratory Syndrome, or MERS), and SARS-CoV (the beta coronavirus that causes severe acute respiratory syndrome, or SARS) (Su et al., 2016).

### Sequence featurization and modelling

RNA-GPS uses k-mer featurization with k = 3, 4, 5, applied independently to the 5’ untranslated region (UTR), coding sequence (CDS), and 3’ UTR parts of the transcript (Wu et al., 2020b). Extending this definition to the coronavirus sgRNA sequences, we consider the shared 5’ leader sequence the fixed 5’ UTR input to our model, shared 3’ UTR sequence the fixed 3’ UTR input to our model, and the variable sgRNA sequence the “CDS” input. For sake of consistency with sgRNA transcript mechanisms, this “CDS” sequence includes the current reading frame, along with any 3’ downstream bases until the shared 3’ UTR region begins. Each sgRNA is individually assigned predicted localizations.

For the deep recurrent model, we implemented and trained a recurrent neural network that consumes raw bases as input, maps these to a 32-dimensional embedding layer, passes these through two 64-dimensional gated recurrent units (GRU), and finally a fully-connected layer with sigmoid activation producing 8 localization predictions. This flavor of GRU network is popular in sequence modelling and uses “gating” mechanisms to enable improved learning of longer-range sequence dependencies (Chung et al., 2014). The model was trained using the Adam optimizer (Kingma and Ba, 2014) with a batch size of 1, and early stopping based on validation set AUROC.

Both RNA-GPS and the GRU model are trained and tuned on the same APEX-seq data, measuring localization within HEK293T cells (Fazal et al., 2019), using identical data splits of 80% train, 10% validation, and 10% train. Please see our RNA-GPS manuscript for additional information regarding dataset and modelling details (Wu et al., 2020b).

### Baseline construction and rank score

Baseline distributions are constructed by running a set of baseline transcript sequences through a model predicting transcript localization. For both models (RNA-GPS and the auxiliary GRU model), there is a per-localization baseline derived from human APEX-seq measurements, and one derived from human coronaviruses excluding SARS-CoV-2. For each localization compartment within the human baseline, we consider only transcripts that exhibit significant localization to that compartment, as defined by having a logFC > 0 and adjusted p-value ≤ 0.05 when running differential expression analysis against the remainder of the cell. Additionally, we only use transcripts not used for model training/tuning (i.e. the test data split), as this most closely approximates what the model would predict when presented with novel sequences. For the coronavirus baseline, we do not have systematically measured localization data, so we do not filter as such. However, we make slight adjustments to the process of calculating the rank score (see below). Note that due to these differences, the values produced by these two baselines are not directly comparable.

For these baselines, we define a rank score as the proportion of baseline values that a SARS-CoV-2 sgRNA localization prediction exceeds. A hypothetical value of 0.5 would correspond to a median, 0.25 would correspond to the first quartile, etc.; rank score is thus bound between 0 and 1. Note that this rank is calculated for each individual compartment separately, as the baselines themselves are compartment specific. For the coronavirus baseline, each SARS-CoV-2 sgRNA is also compared only to homologous sgRNAs from other coronaviruses. For example, the spike protein’s localization prediction is only compared against localization predictions of other coronavirus spike proteins. This limits our comparison to the set of genes with easily traceable homology across human coronaviruses, namely ORF1ab, spike (S), envelope (E), membrane (M), and nucleocapsid (N) (Woo et al., 2010). As previously discussed, localization predictions are averaged across all valid SARS-CoV-2 genomes prior to calculating rank scores.

### RNA binding protein PWM identification and ablation

We use a database of 102 RNA binding protein binding motifs (Ray et al., 2013). To identify matches, we use the same methodology as was used in the RNA-GPS manuscript (Wu et al., 2020b). Briefly, we start with the position weight matrix (PWM) that describes the motif, adjust its probabilities to account for the background nucleotide composition of each sequence, define a cutoff score slightly lower than the maximum achievable log-likelihood for that PWM, and identify any subsequences that exceed that cutoff (see RNA-GPS manuscript for additional details).

When ablating these PWMs, we use the same methodology for identifying hits, and subsequently replace all hits with “N” bases, re-featurizing the ablated sequence as necessary before feeding into the model, thus generating the ablated localization predictions.

### Plotting and additional metrics

All plots were generated using a combination of seaborn and matplotlib Python packages (Hunter, 2007). Statistical testing was done using functions available within the scipy Python package (Virtanen et al., 2020), and multiple hypothesis testing correction was done using the statsmodels Python package (Seabold and Perktold, 2010).

## Supplementary Figures

**Supplementary Figure 1:**
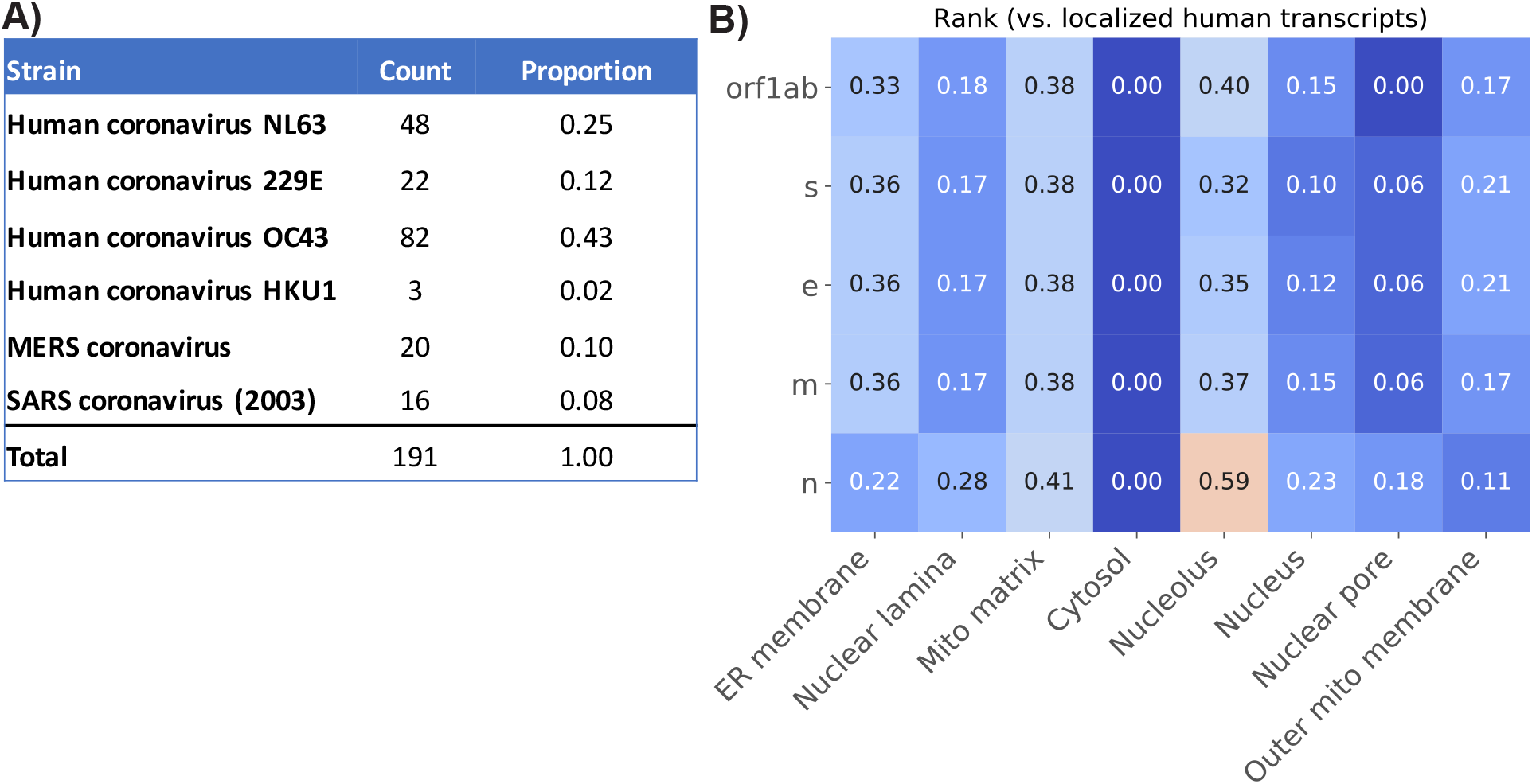
Summary of the human coronavirus baseline. (a) shows the different viral strains comprising the human coronavirus baseline. (b) shows the localization patterns aggregated across all transcripts comprising the human coronavirus baseline. We see that coronaviruses in general primarily exhibit localization towards the nucleolus, mitochondrial matrix, and ER membrane – a pattern similar to that seen in SARS-CoV-2 (albeit less dramatic).

**Supplementary Figure 2:**
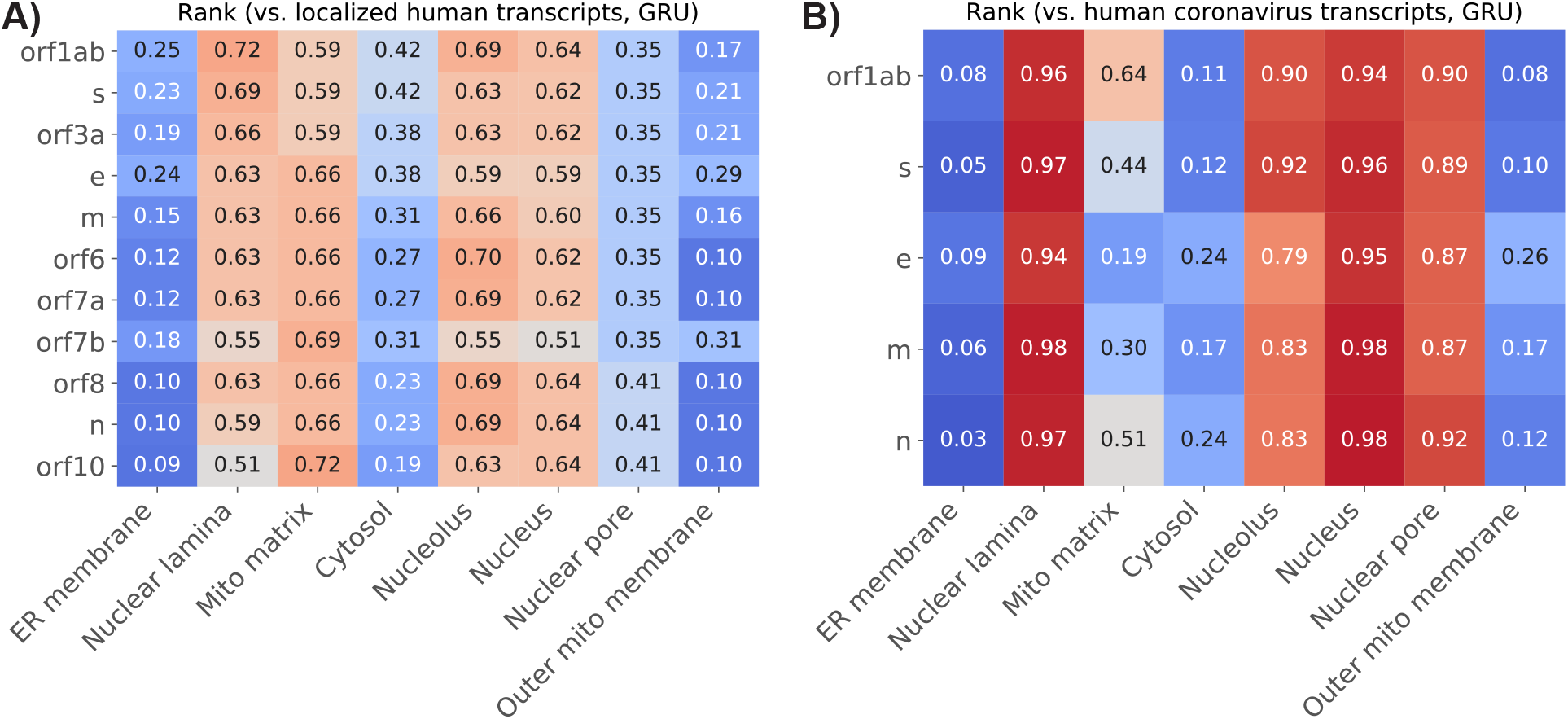
Heatmaps of rank scores of SARS-CoV-2 localization predictions, relative to localized human transcripts (a) and other coronavirus genomes (b), according to a deep-learning recurrent model (GRU). This model takes a very different computational approach to predicting localization, and thus serves as an “orthogonal” validation of results covered in our primary figures. (a) Recapitulates that mitochondrial matrix and nucleolus are among the two most prominent localization signals for SARS-CoV-2. (b) Recapitulates that compared to other coronaviruses, SARS-CoV-2 generally exhibits a stronger nuclear localization signal.

**Supplementary Figure 3:**
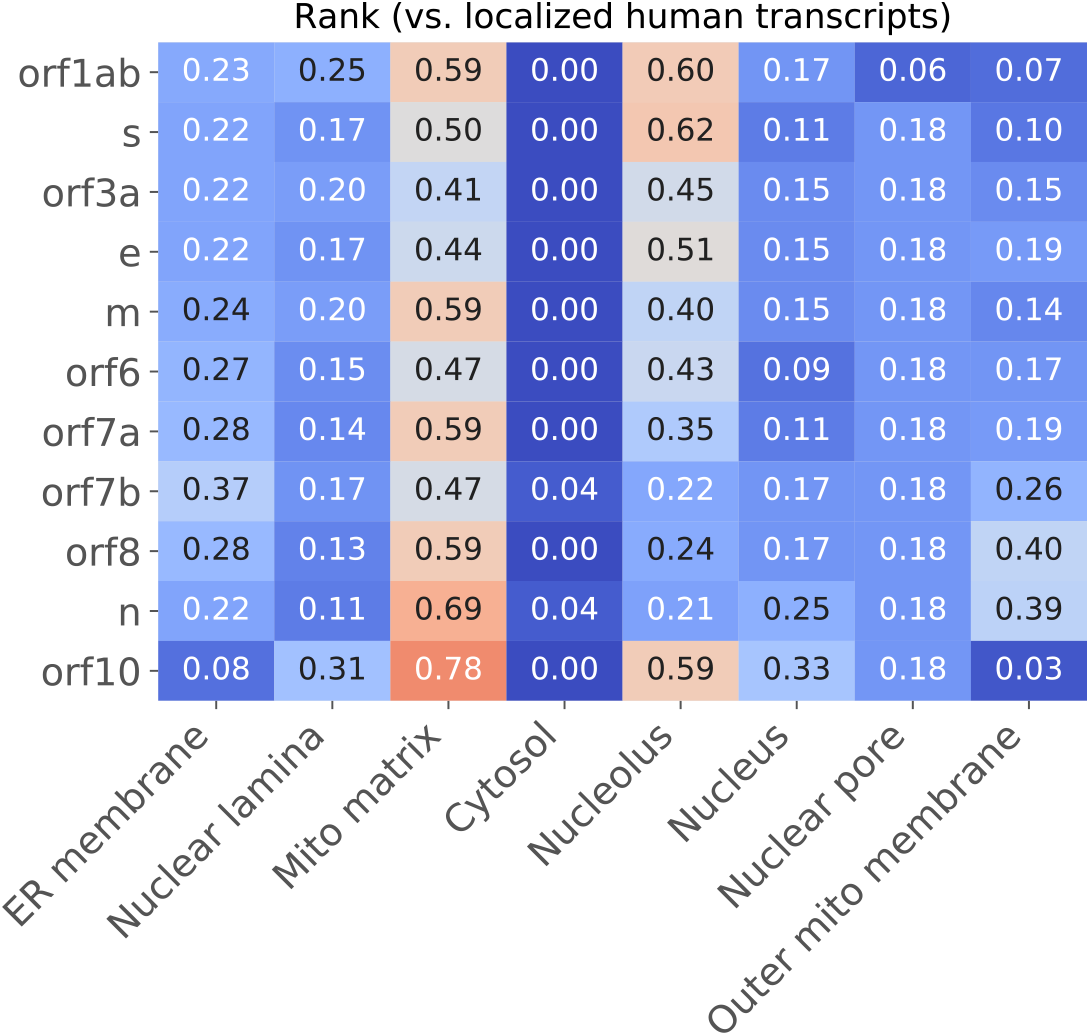
Localization of negative strand sgRNA precursors. Figure 1c shows that the positive strand sgRNA transcripts tend to exhibit localization towards the mitochondrial matrix and nucleolus. Here, we look at the negative-strand precursors to those sgRNAs and observe that these transcripts share similar mitochondrial matrix and nucleolus localization patterns. This is consistent with literature studying co-localization of coronavirus double-stranded RNA and suggests yet another layer of conservation of this localization signal.

